# Conserved allosteric pathways for activation of TRPV3 revealed through engineering vanilloid-sensitivity

**DOI:** 10.1101/443689

**Authors:** Feng Zhang, Kenton Swartz, Andres Jara-Oseguera

## Abstract

The Transient Receptor Potential Vanilloid 1 (TRPV) channel is activated by an array of stimuli, including heat and vanilloid compounds. The TRPV1 homologues TRPV2 and TRPV3 are also activated by heat, but sensitivity to vanilloids and many other agonists is not conserved among TRPV subfamily members. It was recently discovered that four mutations in TRPV2 are sufficient to render the channel sensitive to the TRPV1-specific vanilloid agonist resiniferatoxin (RTx). Here we show that mutation of six residues in TRPV3 corresponding to the vanilloid site in TRPV1 is sufficient to engineer RTx binding. However, robust activation of TRPV3 by RTx requires facilitation of channel opening by introducing individual mutations in the pore, temperatures > 30°C, or co-stimulation with another agonist. Our results demonstrate that the energetics of channel activation can determine the apparent sensitivity to a stimulus and suggest that allosteric pathways for activation are conserved in the TRPV family.

## Introduction

Transient receptor potential (TRP) cation channels are involved in a diverse array of physiological functions (Li, 2017), with many of them acting as sensory detectors of stimuli such as temperature, natural products and various cell-signaling molecules (Flockerzi, 2007). Structurally, TRP channels are tetramers, with each subunit containing six transmembrane helices (S1-S6); the S5 through S6 helices form the central ion-conducting pore, and the S1-S4 helices of each subunit form voltage sensor-like domains located peripherally in a domain-swapped arrangement (Madej and Ziegler, 2018).

Consistent with the remarkable diversity of biological roles that TRP channels play, the types of stimuli that activate these channels also largely differ between TRP subtypes, even between closely related members of the same TRP subfamily (Flockerzi, 2007; Li, 2017). However, many important structural features are highly conserved within each TRP subfamily, and TRPs in general (Palovcak et al., 2015; Bae et al., 2018; Kasimova et al., 2018a; Madej and Ziegler, 2018; McGoldrick et al., 2018; Zheng et al., 2018a; Zheng et al., 2018b; Zubcevic et al., 2018a), raising the possibility that TRPs that are sensitive to different stimuli share mechanisms of activation downstream of their ligand-interaction sites. This is well exemplified by TRPV1 and its closest homologue, TRPV2 (48.4% of sequence identity). TRPV1, an integrator of pain-producing stimuli in nociceptors(Moore et al., 2018), is the only member of the TRPV (vanilloid) subfamily of TRP channels that can be activated by vanilloid compounds (Caterina et al., 1997; Jordt and Julius, 2002), cysteine-reactive molecules (Salazar et al., 2008), and some agonists that directly interact with the extracellular face of the pore domain, such as protons (Tominaga et al., 1998; Jordt et al., 2000) and the double-knot toxin (DkTx) from tarantula venom (Bohlen et al., 2010; Cao et al., 2013; Gao et al., 2016). In contrast, TRPV2 is activated by very few known stimuli, including heat (Caterina et al., 1999; Yao et al., 2011), the non-selective channel modulator 2-aminoethoxydiphenyl borate (2-APB) (Hu et al., 2004) and cannabinoid compounds (Qin et al., 2008), all of which also influence the activity of TRPV1 (Hu et al., 2004; Qin et al., 2008). In this context, it is remarkable that only four mutations in TRPV2 (TRPV2-4M) are sufficient to enable robust activation by the TRPV1-specific vanilloid agonist resiniferatoxin (RTx) (Yang et al., 2016; Zhang et al., 2016), suggesting that the sites where vanilloids interact with TRPV1 and TRPV2-4M are very similar, and that these channels share mechanisms of activation. Indeed, structural studies have confirmed that RTx binds to the same site between the S1-S4 domains and the pore domain in both TRPV1 (Cao et al., 2013; Gao et al., 2016) and TRPV2-4M (Zubcevic et al., 2018b) (Fig. 1 Supplement 1A).

**Figure 1.**
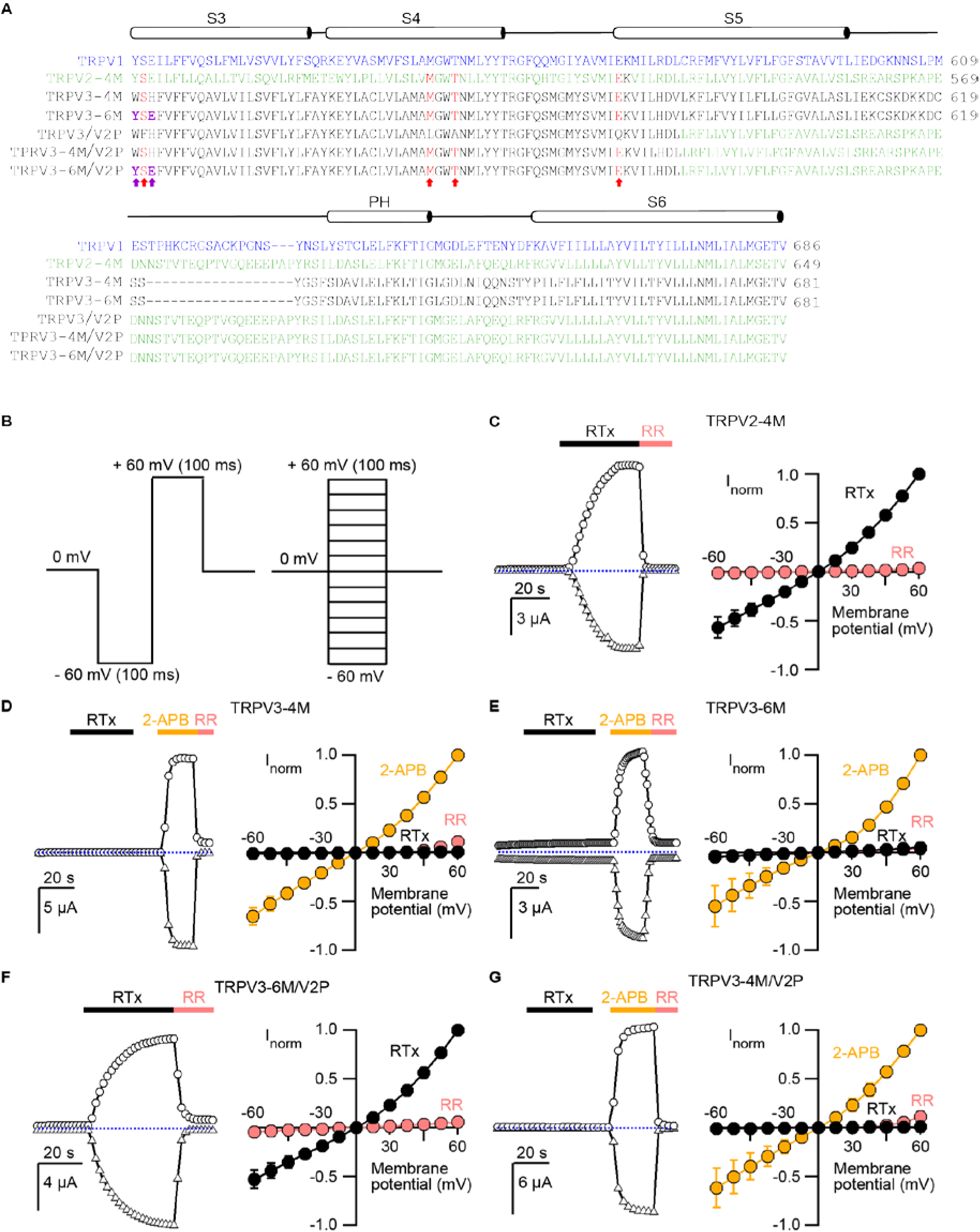
Engineering an RTx binding site into the TRPV3 channel. **(A)** Sequence alignment of S3 through S6 TM helices of rat TRPV1 (blue), rat TRPV2-4M (F472S /L507M/S510T/Q530E, green), mouse TRPV3-4M (F522S/L557M/S560T/Q580E, black), mouse TRPV3-6M (W521Y/H523E/F522S/L557M/A560T/Q580E, black) and mouse TRPV3 chimeras containing the pore domain of rat TRPV2 (TRPV3/V2P). Key residues involved in RTx binding in TRPV2-4M and TRPV3-4M that were mutated to the corresponding residues in TRPV1 are shown in red, with the two additional mutations (W521Y/H523E) in TRPV3-6M shown in purple and bold. **(B)** Voltage protocols used to measure time courses for activation (left) or I-V relations (right). In both cases voltage steps were elicited every 2 s. **(C-G, left panel)** Representative time courses of activation in response to RTx (100 nM) or 2-APB (3 mM) measured at ±60 mV. 50 μM ruthenium red (RR) was applied at the end of each recording to inhibit the channel. The dotted horizontal line indicates the zero-current level. The thick colored horizontal lines indicate the application of agonists or RR. **(C-G, right panel)** Mean normalized I-V relations obtained in the presence of 100 nM RTx (black), 3 mM 2-APB (yellow) and 50 μM RR (red filled symbols). Currents were normalized to the current value in the presence of a saturating concentration of 2-APB (3 mM) or RTx (100 nM) at +60 mV. Data are expressed as mean ± S.E.M (n = 4-8 cells).

In the present study we explored whether vanilloid sensitivity could be engineered into the TRPV3 channel, which shares 42% sequence identity with TRPV1. TRPV3 is expressed primarily in mammalian keratinocytes, where it is required for the formation of the skin barrier (Cheng et al., 2010), and mutations in the channel can lead to skin diseases such as Olmsted Syndrome (Lin et al., 2012; Zhi et al., 2016). TRPV3 is a polymodal detector of warm temperatures (Peier et al., 2002; Smith et al., 2002; Xu et al., 2002), chemicals such as 2-APB (Chung et al., 2004; Hu et al., 2004) and multiple natural products such as camphor, eugenol and thymol (Xu et al., 2006; Vogt-Eisele et al., 2007). To probe for RTx sensitivity in TRPV3, we mutated 6 residues in TRPV3 (TRPV3-6M) corresponding to the vanilloid binding pocket of TRPV1, and found that although RTx can bind to the mutant channel, activation of TRPV3-6M by RTx requires facilitation of the opening transition, either by introducing mutations in the pore domain, or sensitizing the channel with heat or 2-APB. Our results provide functional evidence for conserved mechanisms of activation within the TRPV family, while also providing important information on TRPV3-specific activation properties.

## Results

### Engineering vanilloid sensitivity into the TRPV3 channel

We began by testing whether RTx sensitivity could be engineered into the TRPV3 channel. The vanilloid-binding pocket region is highly similar in TRPV1, TRPV2 and TRPV3, both in terms of amino acid sequence conservation (Fig. 1A) and 3-dimensional structure (Cao et al., 2013; Gao et al., 2016; Huynh et al., 2016; Zubcevic et al., 2016; Singh et al., 2018; Zubcevic et al., 2018b) (Fig. 1 Supplement 1A), and four mutations are sufficient to confer RTx sensitivity to the rat and mouse TRPV2 channels (Yang et al., 2016; Zhang et al., 2016). We initially introduced four mutations into TRPV3 that are equivalent to those in TRPV2-4M to generate the TRPV3-4M channel (F522S/L557M/A560T/Q580E; Fig. 1A and Fig. 1 Supplement 1A), expressed the construct in *Xenopus laevis* oocytes and used the two-electrode voltage-clamp technique to test for responses to RTx and to the non-specific TRPV channel agonist 2-APB (Hu et al., 2004). We recorded current time courses for channel activation at room temperature by each agonist (see Fig. 1B, left panel and Methods), and obtained current-voltage (I-V) relations in the absence and presence of activators, or in the presence of ruthenium red (RR), a non-specific inhibitor of TRP channels (see Fig. 1B, right panel and Methods). Whereas TRPV2-4M displayed robust responses to RTx (Fig. 1C), as we showed previously (Zhang et al., 2016), TRPV3-4M was activated by 2-APB but not by 100 nM RTx (Fig. 1D), a concentration that is saturating for both TRPV1 and TRPV2-4M (Zhang et al., 2016). We found two additional positions within the vanilloid pocket where the size, shape and polarity of the residues in TRPV3 largely differ from those in TRPV1 (H523 and W521 in TRPV3 correspond to E513 and Y511 in TRPV1, respectively; Fig. 1A and Fig. 1 Supplement 1A). We therefore mutated these two additional positions in TRPV3-4M to the corresponding residues in TRPV1 to generate TRPV3-6M, and found that the construct remained insensitive to RTx, although it retained robust responses to 2-APB (Fig. 1E).

The binding of an agonist to its site requires functional coupling to the pore domain to promote opening of the ion conduction pathway. To test whether differences in the pore domain might play a role in preventing RTx activation in TRPV3-6M, we transferred the pore domain of TRPV2 into the TRPV3-6M construct to generate TRPV3-6M/V2P (Fig. 1A). Remarkably, the resulting chimera was robustly activated by RTx and readily blocked by RR (Fig. 1F), indicating that the pore domain of TRPV2 contains determinants that are important for RTx-activation. We also found that the two additional residues in the vanilloid pocket differing between TRPV3-4M and TRPV3-6M were necessary for RTx activation, since we transferred the pore domain of TRPV2 into TRPV3-4M to generate TRPV3-4M/V2P (Fig.1A), and found that the resulting chimera could be activated by 2-APB but not by RTx (Fig. 1G).

### Mutations within the pore domain required for vanilloid activation of TRPV3

We next sought to identify the specific residues within the pore domain of TRPV2 that enable activation of TRPV3-6M/V2P by RTx. We aligned the sequences of TRPV1, TRPV2 and TRPV3 and found 21 amino acids in the pore domain that are conserved between TRPV1 and TRPV2 but differ in TRPV3 (Fig 2A). We individually mutated each of the 21 residues in the TRPV3-6M background to those in TRPV1 and TRPV2, and tested for their response to RTx and 2-APB. We identified five positions that resulted in robust activation by RTx (Fig 2B, C). Notably, these five residues are widely distributed within the pore domain, located in the S5 helix (V587L and A606V), pore turret (F625L), and S6 helix (F656I and F666Y) (Fig. 2D). In addition, five other mutants exhibited detectable responses to RTx (I595L, K611D, L635F, L639M, Y650F; Fig 2B-D). Interestingly, some of the mutations that conferred RTx sensitivity to TRPV3-6M are very similar to WT amino acids, such as V587L and F666Y, whereas some mutations expected to be more disruptive, such as D618P, S620E or P651K, did not have an apparent effect on RTx- or 2-APB-senstivity (Fig. 2B, C). Given their considerable distance from the vanilloid binding pocket (Fig. 2D), it seems unlikely that any of the residues that had large effects on the response of TRPV3-6M to RTx directly interact with the vanilloid. The observation that each of the 10 identified residues enable RTx activation when mutated individually, together with their wide-spread distribution within the pore domain, support the idea that allosteric interactions influencing activation can be delocalized in TRPs (Clapham and Miller, 2011).

**Figure 2.**
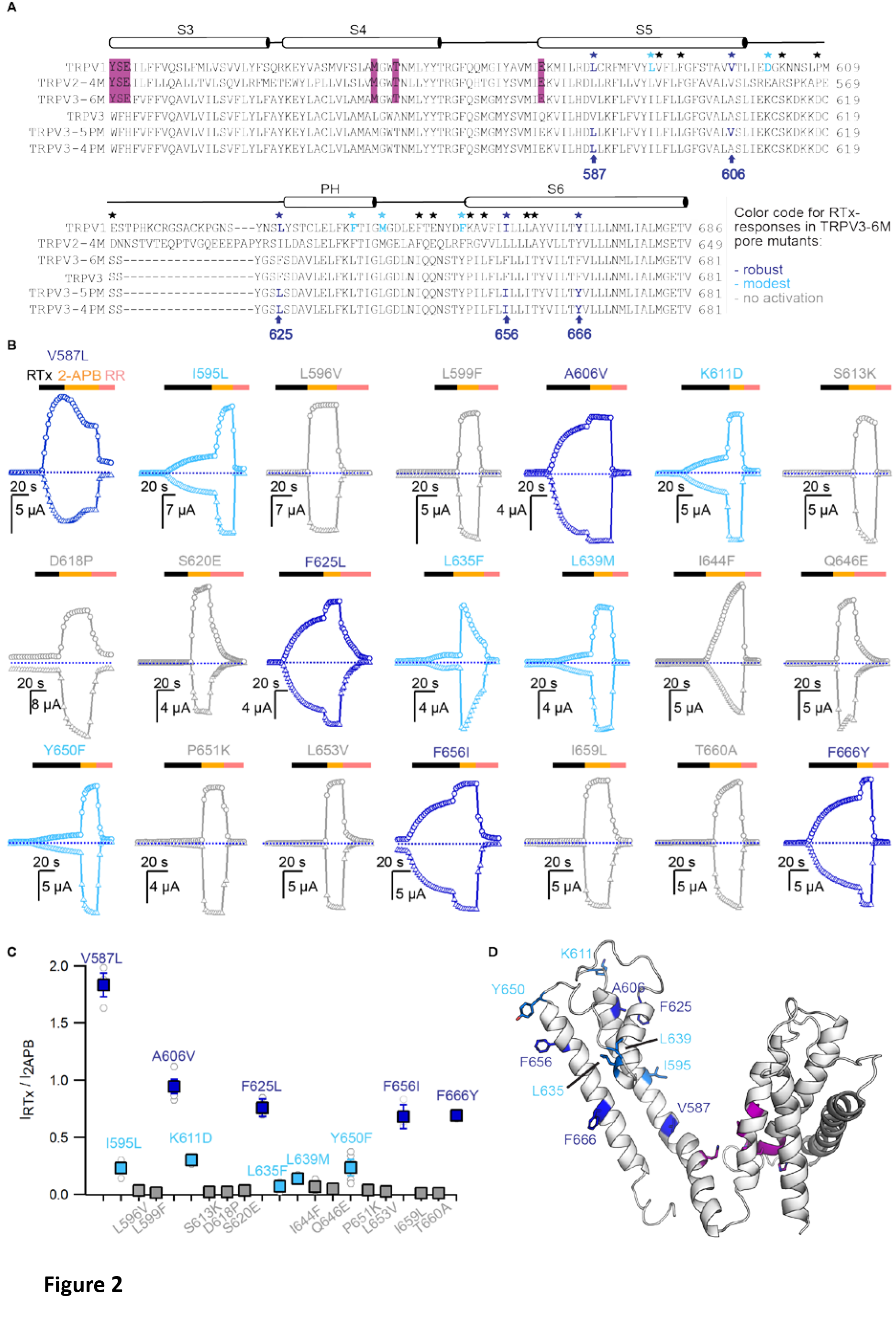
Identification of key residues for RTx activation in the pore domain of TRPV3-6M. **(A)** Sequence alignment of the S3 to S6 TM helices for rTRPV1, rTRPV2-4M, mTRPV3-6M, WT mTRPV3, TRPV3-5M and TRPV3-4M. Residues labeled with stars are conserved in TRPV1 and TRPV2 but different in TRPV3; black stars – mutations that did not influence RTx activation; blue - mutations that enabled moderate (light blue) or strong (dark blue) responses to RTx. The purple highlights denote the 6M mutations. **(B)** Representative time courses of activation of TRPV3-6M channels with individual pore mutations. Channels were stimulated by RTx (100 nM) and 2-APB (3 mM), and blocked with RR (50 μM) as indicated by the colored horizontal lines. Currents were measured at +60 (circles) and -60 mV (triangles) as in Fig. 1. The dotted horizontal lines indicate the zero-current level. **(C)** Summary of the current magnitudes activated in response to RTx relative to saturating 2-APB at +60 mV from experiments as in (B). Values for individual oocytes are shown as open circles and mean ± S.E.M. as squares (n = 3-6). **(D**) Side view of a cartoon representation of the transmembrane domain of a mTRPV3 subunit (PDB ID code: 6DVW)(Singh et al., 2018). Side-chains in dark blue are from the residues that enable strong RTx activation when mutated. Light blue is for residues that enable weak RTx activation. Residues within the RTx binding pocket that were mutated in the 6M construct are highlighted in purple.

### Temperature sensitivity of TRPV3 constructs

The binding of each agonist to its site in the channel must allosterically influence the conformation of the pore domain to open the ion conduction pathway. Therefore, it is possible that the mutations in the pore that enable RTx-dependent activation of TRPV3 do so by shifting the gating equilibrium towards the open state, which would facilitate activation by RTx and other stimuli as well. Indeed, opening of TRPV3 upon initial stimulation seems to be largely disfavored when compared to TRPV1 and TRPV2 (Liu et al., 2011; Liu and Qin, 2016). For example, the TRPV3 channel can only initially be activated by heat when temperatures > 50°C are applied, but will respond to much lower temperatures on subsequent heating (Liu et al., 2011). To test if the five pore mutations affect the overall gating equilibrium, we determined if they facilitate TRPV3 activation by heat. For these experiments we used the two-electrode voltage clamp technique together with a fast temperature-controlled perfusion system, as previously described (Fig. 3A) (Zhang et al., 2018). First, we detected robust activation by heat in TRPV1-expressing oocytes, which we used as a positive control (Fig. 3B). For WT TRPV3, we failed to observe any response to a single heat stimulus < 50°C, but observed robust responses to 2-APB (Fig. 3C, F), consistent with previous reports (Liu et al., 2011; Liu and Qin, 2017). We observed the same result for TRPV3-6M, indicating that the 6M mutations do not noticeably influence channel activation or temperature-sensing (Fig. 3 Supplement 1A). We then measured the responses to heat and 2-APB of five TRPV3 constructs without the 6M mutations, each containing one of the pore mutants that gave rise to robust RTx activation in TRPV3-6M. Four out of five single point mutations (V587L, F625L, F656I and F666Y) displayed strong responses to 2-APB but no activation by temperatures below 50°C (Fig. 3F and Fig. 3 Supplement 1 B-E). In contrast, the A606V mutant exhibited temperature-activated currents below 45°C, which were clearly larger than the responses of WT TRPV3 and TRPV3-6M, but slightly smaller than heat-driven activation in TRPV1 when compared to responses to 2-APB (Fig. 3 D, F). This result suggests that the mutation at A606 facilitates TRPV3 activation through multiple modalities, including RTx and heat. We also found that the responses to heat of the TRPV3/V2P chimera without the 6M mutations (Fig. 3 Supplement 1F), and the TRPV3-5PM construct with all five cumulative pore mutations in the WT TRPV3 background (i.e. without the 6M mutations) (Fig.3 E, F) were qualitatively similar to that of the A606V mutant, suggesting that the other four positions in the pore do not contribute to temperature-dependent activation. Consistently, a construct containing all pore mutations except A606V (TRPV3-4PM) exhibited no response to temperatures below 50°C (Fig. 3 Supplement 1G).

**Figure 3.**
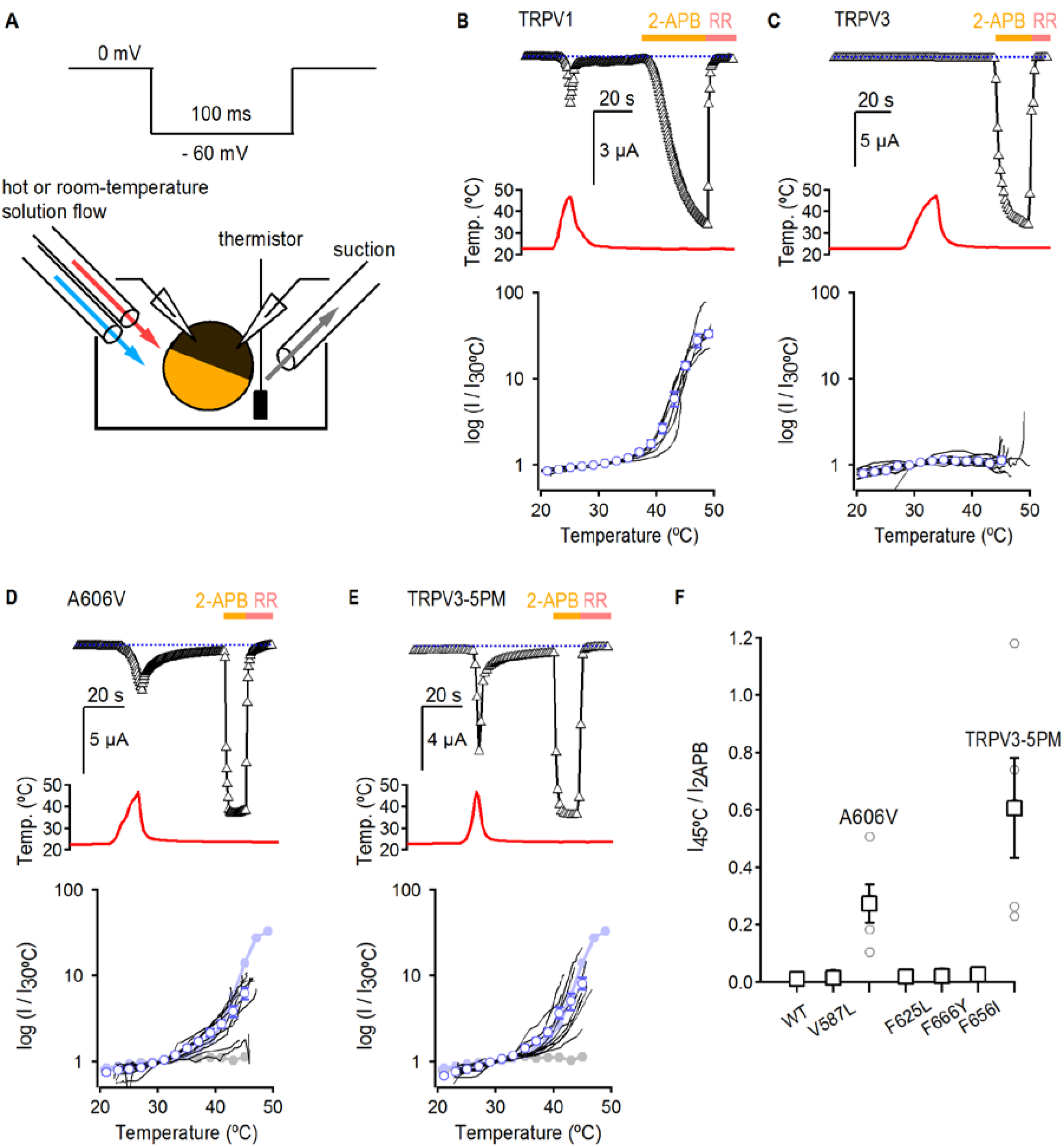
Temperature sensitivity of TRPV3 constructs. **(A)** Voltage protocol used for recording the temperature responses (top), and a cartoon of the temperature-control system used for the experiments (bottom). Temperature was controlled by using two perfusion lines immersed in baths at high-or room-temperature (Zhang et al., 2018), and temperature was measured with a thermistor positioned close to the oocyte. **(B)** Representative time-course obtained from a TRPV1-expressing oocyte (upper panel), showing the response to a heating stimulus, followed by the application of 3 mM 2-APB and RR (50 μM). The dotted line indicates the zero-current level. The recorded temperature is shown in the middle panel. The bottom panel shows the logarithm of the currents at -60 mV normalized to their value at 30°C plotted as a function of temperature, obtained from experiments as in the upper panel. Data from individual cells are shown as black curves, and the mean ± S.E.M as blue open circles (n=7). **(C-E)** Representative current-(top panel) and temperature-(middle panel) time courses obtained from the constructs indicated. The lower panel shows the log(I)-temperature relations obtained from experiments as in the upper panel (n=7–15). In (D) and (E), the curves in solid circles correspond to the mean log(I) vs T relations for WT TRPV1 (light blue) and TRPV3 (grey). **(F)** Summary of current responses to heat (46°C) relative to saturating 2-APB (3 mM) at room temperature. Values for individual oocytes are shown as open circles and mean ± S.E.M. as open squares (n = 4-5).

### Residues in the pore have distinct effects on activation by different stimuli

The results thus far indicate that A606V facilitates activation by temperature and enables RTx sensitivity, whereas the other four mutated positions in the pore domain have no effect on temperature-dependent activation. We wondered whether these four mutations might favor activation by stimuli other than RTx, such as 2-APB. We therefore set out to measure the effects of pore mutations introduced into the WT background (i.e. no 6M mutations) on concentration-response relations for activation by 2-APB. We first measured the response of WT TRPV3 to 2-APB, and obtained an apparent affinity of 460 ± 12 μM (Fig. 4A, F), consistent with a previous report (Phelps et al., 2010). Interestingly, the V587L pore mutant, which robustly promotes activation by RTx (Fig. 2B), did not have any effect on the apparent affinity for 2-APB (440 ± 13 μM; Fig. 4B, F). In contrast, the A606V mutation produced a detectable (~3.5-fold) increase in the apparent affinity to 132 ± 7 μM (Fig. 4C, F), consistent with this mutation having a generalized effect on TRPV3 channel gating, as it affects the sensitivity of the channel to all tested stimuli (Liu et al., 2011). Interestingly, we found that a construct containing four cumulative pore mutations without A606V (TRPV3-4PM) exhibited a ^~^2-fold shift in the apparent affinity for 2-APB (210 ± 2 μM, Fig. 4D, F), which combined synergistically with A606V in TRPV3-5PM for a 20-fold increase in the apparent affinity (19 ± 2 μM, Fig. 4E, F). Together, these results suggest that the five residues in the pore facilitate opening by distinct mechanisms, differentially affecting activation by heat and 2-APB, while each enabling activation by RTx. Notably, the maximal response to 2-APB was similar to the response to RTx in the 4PM and 5PM mutant channels when tested in the 6M background (TRPV3-6M/4PM and TRPV3-6M/5PM) (Fig. 4 Supplement 1), as well as in all five individual pore mutants when in the 6M background (Fig. 2B, C).

**Figure 4.**
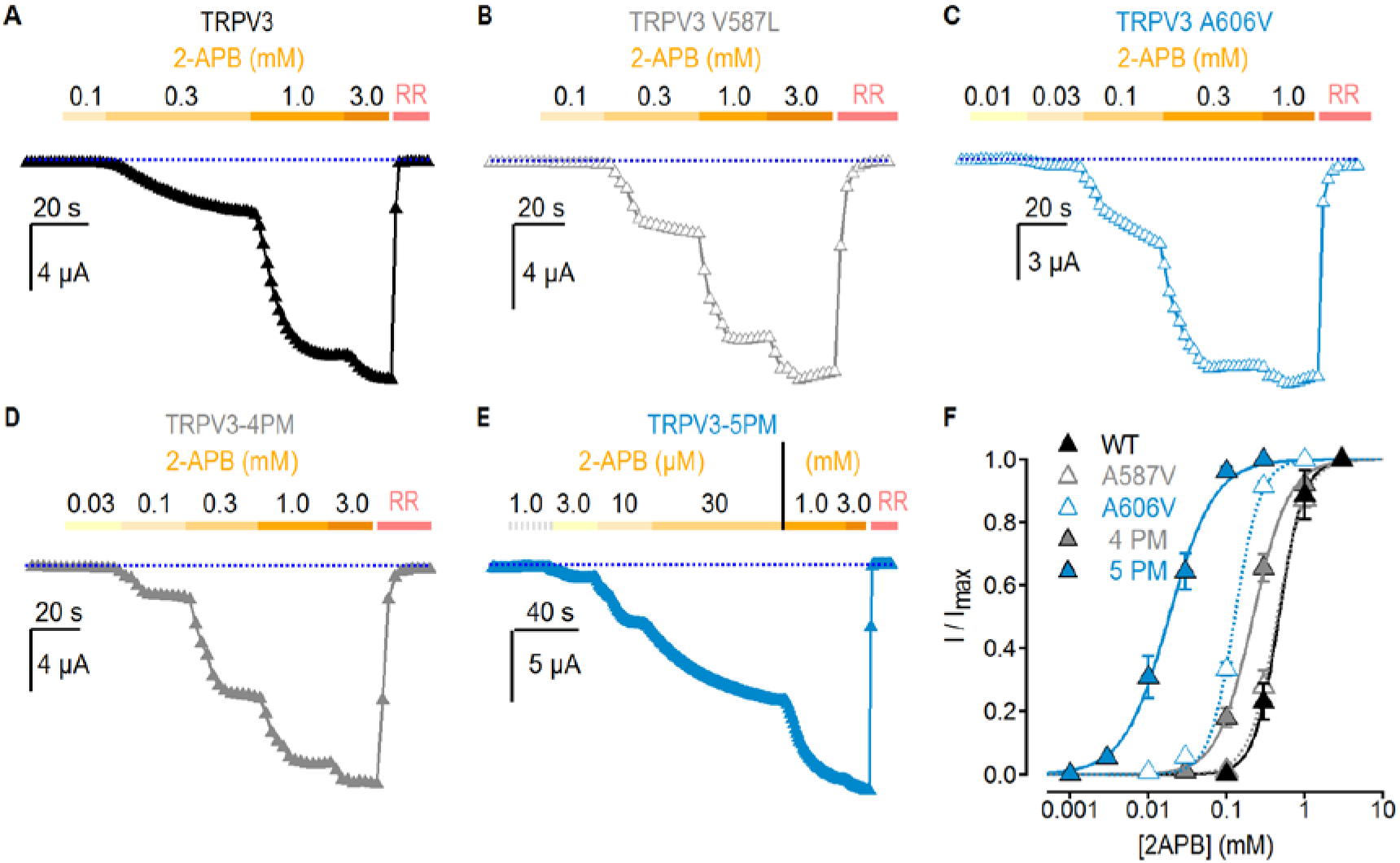
Effect of pore mutations on TRPV3 2-APB sensitivity. (A-E) Representative current time courses at -60 mV at increasing concentrations of 2-APB, followed by block with RR (50 μM), as indicated by the colored horizontal lines. The blue dotted lines indicate the zero-current level. Currents were held at 0 mV and voltage was stepped to -60 mV for 100 ms every 2s. **(F)** Mean normalized concentration-response relations for 2-APB measured from experiments as in (A-E). Data are shown as mean ± S.E.M. (n= 4-6). The continuous curves are fits to the Hill equation with EC_50_ and s (slope) values as follows: TRPV3, EC_50_ = 464 ± 12 μM, s =2.63; TRPV3-4PM, EC_50_ = 210 ± 20 μM, s = 1.84; TRPV3-A606V, EC_50_ = 132 ± 7, s = 2.72; TRPV3-5PM, EC_50_ = 19 ± 2 μM, s = 1.32; TRPV3-V587L, EC_50_ = 443 ± 13 μM, s = 2.28.

### Heat and 2-APB render TRPV3-6M channels sensitive to RTx in the absence of mutations in the pore domain

The results thus far suggest that the A606V mutation favors channel opening in a stimulus-independent manner, and that this is sufficient to allow TRPV3-6M channels to open in response to RTx at room temperature. If this were true, the TRPV3-6M channel should be able to respond to RTx if opening is facilitated by other stimuli that activate TRPV3, such as heat or 2-APB. Consistent with the WT TRPV3 channel being unable to bind RTx, as it lacks the 6M mutations in the binding pocket, application of the vanilloid together with heating did not elicit any response (Fig. 5A). Consistent with our hypothesis, application of a short heating pulse together with RTx resulted in prominent activation of TRPV3-6M at temperatures only slightly above room temperature (Fig. 5B), in contrast to the lack of response to heating up to 45°C in the absence of RTx for the same construct (Fig. 3 Supplement 1A). These results indicate that RTx and temperature act cooperatively to activate TRPV3-6M under conditions where neither stimulus alone would suffice. Consistent with cooperativity between heat and RTx binding, and our proposed mechanism for A606V, both the TRPV3-6M/V2P chimera and TRPV3-6M A606V responded to RTx at room temperature (Figs. 1F and 2B), and TRPV3 A606V was activated by a heating stimulus < 50 °C (Fig. 3E). In addition, activation of the TRPV3-6M/V2P chimera to RTx observed at room temperature could be reduced to baseline levels when temperature was decreased to 10°C (Fig. 5C), providing further evidence of coupling between heat-dependent activation and RTx sensitivity. We also examined whether we could sensitize RTx activation of TRPV3-6M by pre-stimulation with a saturating concentration of 2-APB. Similar to our results with heat, application of RTx after a short stimulation with 2-APB did not activate WT TRPV3 channels (Fig. 5D), but resulted in robust activation of TRPV3-6M channels in the absence on additional pore mutations (Fig. 5E). Collectively, these results demonstrate that RTx binds to the TRPV3-6M channel, but that further facilitation of channel opening is required for the vanilloid to activate TRPV3-6M.

**Figure 5.**
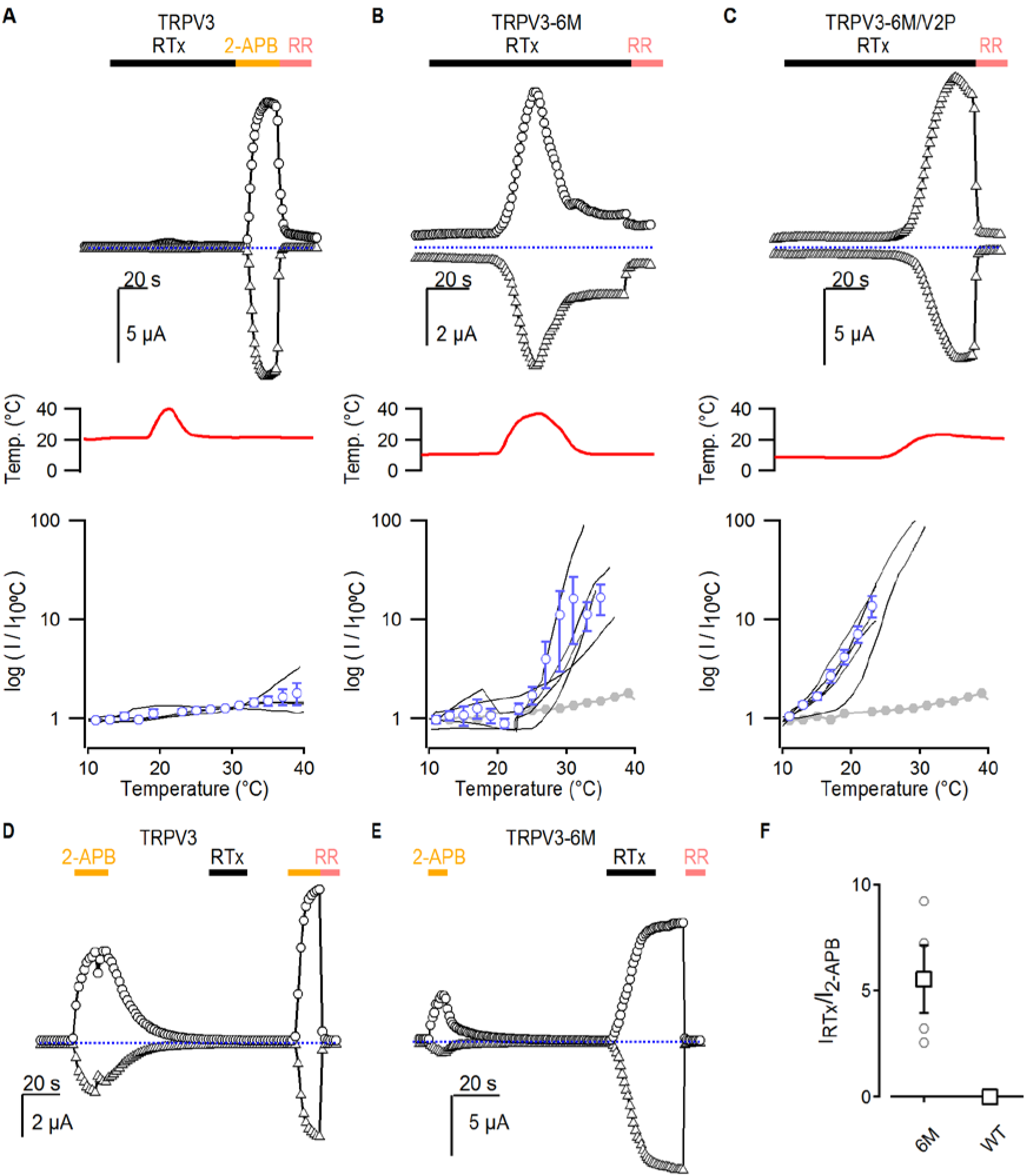
Interaction between RTx, temperature and 2-APB in TRPV3 channels. (A-C) Representative time courses for (A) WT TRPV3, (B) TRPV3-6M and (C) TRPV3-6M/V2P activation in response to 100 nM RTx and changes in temperature, which is shown in the middle panel. WT TRPV3 was also activated by 2-APB (3 mM), and all currents were blocked by RR (50 μM). Time courses were obtained at +60 (circles) and -60 mV (triangles) as described in Fig. 1B, and temperature was controlled and monitored as in Fig. 3. The blue dotted horizontal lines indicate the zero-current level. The bottom panel shows normalized log(current) vs temperature relations at -60 mV obtained from experiments as in the upper panels. Individual cells are shown as black curves, with currents normalized to their amplitudes at 10 °C, and the mean ± S.E.M. is shown in blue open circles (n = 3-4). For (B) and (C), the mean log(I) vs T for WT TRPV3 is depicted in solid grey circles. **(D, E)** Representative time courses of activation of (D) WT TRPV3 and (E) TRPV3-6M in response to 2-APB (3 mM), RTx (100 nM), and RR (50 μM) at ± 60 mV. **(F)** Summary of the current amplitude ratios for TRPV3 and TRPV3-6M activation by RTx relative to 2-APB. Values for individual oocytes are shown as open circles, the mean ± S.E.M. in open squares (n = 4-6).

## Discussion

The goal of the present study was to determine whether sensitivity to vanilloids could be engineered into the TRPV3 channel. Our results demonstrate that like TRPV2, the TRPV3 channel contains a ‘defunct’ vanilloid binding site that can be readily made functional by mutating a few key positions in the binding pocket. More importantly, our results show that the allosteric machinery required for coupling vanilloid-binding to pore-opening has remained functional through evolution in both TRPV2 and TRPV3 channels, even though these last two channels are not naturally sensitive to vanilloid molecules. This points to the presence of conserved mechanisms of gating among structurally related but functionally distinct TRP channels. In contrast to TRPV2, RTx binding to the engineered vanilloid site in TRPV3 was not sufficient to promote channel activation at room temperature, which required further manipulations: either co-stimulation with heat or 2-APB, or the introduction of single point mutations at one of 10 identified positions throughout the pore that are conserved in TRPV1 and TRPV2 but differ in TRPV3. This indicates that agonist sensitivity in TRP channels is not only fine-tuned by the structure of the agonist binding pocket, but also by the intrinsic energetics of activation that are influenced by multiple sites throughout the pore. Out of the 5 positions in the pore that enabled robust RTx activation in the TRPV3-6M construct, A606 at the extracellular end of the S5 helix had an effect on temperature-activation and moderately increased sensitivity to 2-APB. In contrast, the other four positions had no clear influence on heat-dependent activation and only produced a large (>20-fold) increase in 2-APB sensitivity when introduced cumulatively and together with A606V. These findings point to the presence of non-overlapping allosteric networks in the pore, and likely the rest of the protein, that selectively influence activation by specific stimuli.

It is intriguing that both TRPV2 and TRPV3 can respond to RTx once the vanilloid pocket has been engineered. Structural data suggest that the vanilloid site can also be occupied by lipids in TRPV1 (Gao et al., 2016), TRPV2 (Zubcevic et al., 2016) and TRPV6 (McGoldrick et al., 2018), raising the possibility that the functional coupling between the vanilloid site and the pore is maintained in TRPV2 and TRPV3 because it is important for the modulation of these channels by lipids. The overall fold of the vanilloid site, located at a key interface between the S1-S4 domain and the pore (Jordt and Julius, 2002; Cao et al., 2013; Gao et al., 2016; Zhang et al., 2016; Singh et al., 2018; Zubcevic et al., 2018b), and the relative position of important side chains in the site are highly conserved between TRPV1, TRPV2 and TRPV3 channels (Fig. 1 Supplement 1A). Indeed, the crystal structure of the TRPV2-4M channel solved with RTx bound (Zubcevic et al., 2018b) suggests a similar binding pose for the vanilloid when compared to TRPV1 (Fig. 1 Supplement 1A). However, the RTx-bound TRPV2-4M structure exhibits two-fold symmetry (Zubcevic et al., 2018b) in contrast to the nearly four-fold symmetric structure of the DkTx/RTx-TRPV1 complex (Gao et al., 2016), raising the possibility that the mechanisms of activation by RTx are somewhat different in the two channels. In addition, a π-helical disruption in the network of backbone hydrogen bonds within the pore-lining S6 helix that was proposed to serve as a gating hinge in TRPs (Palovcak et al., 2015; Kasimova et al., 2018b; Kasimova et al., 2018a) is present in all open and closed TRPV1 structures (Cao et al., 2013; Liao et al., 2013; Gao et al., 2016), but is absent in the TRPV2 structures (Zubcevic et al., 2016; Zubcevic et al., 2018b), which all appear to be in non-conducting states. The π-bulge is also absent in the closed mouse TRPV3 structures, although it is present in the open mouse TRPV3 structure (Singh et al., 2018) and the 2-APB-stimulated human TRPV3 (Zubcevic et al., 2018a). At present it remains unclear whether these differences observed between the structures of TRPV1, TRPV2 and TRPV3 reflect actual differences in their activation mechanisms, or an incomplete sampling of the conformational landscape associated with channel activation.

It is intriguing to consider whether the other members of the TRPV family could be engineered to be activated by RTx. The structure of the zebrafish TRPV4 channel exhibits significant differences relative to all other TRPVs, particularly in the orientation of the S1-S4 domains relative to the pore (Deng et al., 2018). This difference results in large structural distortions in the region of TRPV4 corresponding to the vanilloid pocket of TRPV1 (Fig. 1 Supplement 1B), making it unlikely that RTx-sensitivity could be engineered into TRPV4. The structure of the vanilloid pocket is better conserved in the TRPV5 (Hughes et al., 2018) and TRPV6 (McGoldrick et al., 2018) channels (Fig. 1 Supplement 1B), and structural and functional data indicate that the same pocket where vanilloids interact is occupied by a channel-activating lipid in TRPV6 (McGoldrick et al., 2018), or by the inhibitor econazole in TRPV5 (Hughes et al., 2018), raising the possibility that the pocket may be functionally coupled to the pore in these two channels.

It is interesting that in contrast to TRPV1 and TRPV2-4M, binding of RTx to TRPV3-6M does not readily elicit activation at room temperature. The initial activation of TRPV3 seems to involve a transition with a large energy barrier, as either high concentrations of 2-APB or much higher temperatures (> 50 °C) are initially required for robust activation when compared to subsequent stimulations, which elicit maximal activation at lower temperatures (< 40°C) or 2-APB concentrations (Chung et al., 2004; Liu et al., 2011; Liu and Qin, 2017). Activation of TRPV1 and TRPV2 by heat involves significant hysteresis as well, as both the apparent threshold for activation and the steepness of the current-temperature relations are reduced once channels have been activated by heat (Liu and Qin, 2016; Sanchez-Moreno et al., 2018). However, in contrast to TRPV3, TRPV1 channel opening by the vanilloid capsaicin does not involve hysteresis (Liu and Qin, 2016), suggesting that TRPV1 activation by vanilloids does not necessarily require overcoming a large initial energy barrier. Furthermore, initial activation of TRPV1 and TRPV2 with 2-APB sensitizes both channels to subsequent 2-APB applications (Liu and Qin, 2016), but to a lesser degree than in TRPV3 (Chung et al., 2004; Liu et al., 2011) and more importantly, without influencing the response of TRPV1 or TRPV2 to subsequent heat stimuli (Liu and Qin, 2016). Together, these observations raise the possibility that a large energy barrier is an obligatory component of activation for TRPV3 but not for TRPV1 or TRPV2. The origin of this large energy barrier is not known, but it is interesting to note that the open and closed TRPV3 structures exhibit far more drastic differences than those of open and closed TRPV1 (Cao et al., 2013; Gao et al., 2016) and TRPV6 (McGoldrick et al., 2018) structures, including significant changes in the length of the S6 and TRP helices, and a 20° rotation of the rest of the channel relative to the pore domain (Singh et al., 2018). Furthermore, a structure of human TRPV3 has recently been obtained under conditions that favor sensitization, and it also exhibits significant conformational changes in the S6, TRP box and S4-S5-linker helices as compared to the apo closed state (Zubcevic et al., 2018a). It is therefore possible that RTx binding to TRPV3-6M channels is not sufficient to overcome this large barrier, either because the vanilloid site in TRPV3-6M may lack the structural determinants to influence this slow transition, or because of energetically weaker coupling of the RTx site to the gating machinery. It is likely that the binding of RTx to TRPV3-6M and TRPV2-4M and/or its coupling to the pore might not function as efficiently as in TRPV1. Indeed, RTx-sensitive TRPV2 and TRPV3 constructs cannot be activated by capsaicin (Zhang et al., 2016) (Fig. 5 Supplement 1A and data not shown).

If RTx-binding to TRPV3-6M is not sufficient to overcome the initial energy barrier for activation, it is reasonable to expect that stimuli capable of activating WT TRPV3, such as heat or 2-APB, could enable RTx activation, as we show in the present manuscript. What is then the role of the pore mutations in enabling TRPV3-6M channel activation by RTx at room temperature? The A606V mutant facilitates activation by RTx, temperature and 2-APB, suggesting that it facilitates opening by affecting temperature sensing, the efficacy of coupling between the stimulus-sensing domains and the pore, or by directly stabilizing the sensitized or open states. It will be interesting to investigate how A606 and RTx sensitivity relate to position S404 in the membrane-proximal N-terminal domain that has been associated with temperature-dependent sensitization in TRPV3 (Liu and Qin, 2017). The other pore mutations that also enable robust RTx activation is more puzzling, as none of them had large effects on the response of the channel to temperature or 2-APB. It would be interesting to investigate how different residue substitutions at those sites and their possible interaction partners affect other aspects of function in TRPV1, TRPV2 and TRPV3. Given that the mutations seem to have a negligible effect on channel opening by heat or 2-APB, but strongly promote RTx activation, one possible interpretation is that the mutations allow the RTx binding site to allosterically influence the high-energy transition in the activation pathway without directly causing sensitization. Importantly, the five identified pore mutations added cumulatively to TRPV3 did not have an effect on activation by other TRPV1-specific stimuli, such as extracellular protons (Jordt et al., 2000) or double knot toxin (DkTx) (Bohlen et al., 2010; Cao et al., 2013; Gao et al., 2016) (Fig. 5 Supplement 1B, C), consistent with the absence of binding sites for these two modulators in TRPV3.

The fact that conservative side-chain modifications, such as V587L or I595L, can have such a dramatic effect on the response to RTx in TRPV3-6M raises the possibility that other relatively subtle structural alterations, such as those caused by the binding of another ligand, a post-translational modification, or an alteration in the properties of the membrane, could enable responses to channel modulators that would otherwise seem inactive. The picture of TRPV channel gating that emerges from this study is one where fine structural differences dictate the response of a channel to an agonist or combination of agonists. Understanding these differences might be key to unraveling the mechanisms of allosteric signal integration in TRP channels.

## Methods

### Expression of constructs and molecular biology

The rat TRPV1 and TRPV2 channel (Caterina et al., 1997; Caterina et al., 1999) cDNAs were kindly provided by Dr. David Julius (UCSF), whereas mouse TRPV3 (Peier et al., 2002) cDNA was kindly provided by Dr. Feng Qin (SUNY Buffalo). All constructs were sub-cloned into the pGEM-HE vector for expression in oocytes. Chimeras were generated by a three-step overlapping PCR protocol and mutations were introduced using a two-step overlapping PCR mutagenesis technique. The DNA sequence of all constructs and mutants was confirmed by automated DNA sequencing and cRNA was synthesized using the T7/SP6 polymerase (mMessage mMachine kit, Ambion) after linearizing the DNA with the appropriate restriction enzymes.

### Electrophysiological recordings

All channel constructs were expressed in *Xenopus laevis* oocytes and studied using the two-electrode voltage clamp technique (TEVC) following 1–4 days of incubation after cRNA injection. Oocytes were removed surgically and incubated with agitation for 1 hr in a solution containing (in mM) 82.5 NaCl, 2.5 KCl, 1 MgCl_2_, 5 HEPES, pH 7.6 (with NaOH), and collagenase (2 mg/ml; Worthington Biochemical, Lakewood, NJ). Defolliculated oocytes were injected with cRNA for each of the constructs and incubated at 17°C in a solution containing (in mM) 96 NaCl, 2 KCl, 1 MgCl_2_, 1.8 CaCl_2_, 5 HEPES, pH 7.6 (with NaOH), and gentamicin (50 μg/ml, GIBCO-BRL, Gaithersburg, MD). Oocyte membrane voltage was controlled using an OC-725C oocyte clamp (Warner Instruments, Hamden, CT). Data were filtered at 1–3 kHz and digitized at 20 kHz using pClamp software (Molecular Devices, Sunnyvale, CA) and a Digidata 1440A digitizer (Axon Instruments). Microelectrode resistances were 0.1–1 MΩ when filled with 3 M KCl. For recording TRP channel currents, the external recording solution contained (in mM) 100 KCl, 10 HEPES, pH 7.6 (with KOH). Experiments were performed at room temperature (^~^22°C), except as indicated otherwise. Agonists and RR were applied using a gravity-fed perfusion system that resulted in exchange of the 150 μL recording chamber volume within a few seconds. Current time courses were obtained by stepping membrane voltage every 2 s from a holding potential of 0 mV to either -60 or +60 mV for 100 ms, and plotting the mean steady-state current at the end of each voltage step as a function of total recording time. I-V relations were obtained similarly, but voltage was stepped every 2 s to a different value, in 10 mV increments and starting at -60 mV.

For temperature-activation experiments, calcium-activated chloride currents were minimized by using calcium free solution containing 30 μM Caccinh-A01 and recording at negative membrane voltages. Heat stimuli were achieved by passing the external recording solution through glass capillary spirals immersed in a water bath maintained at about 70 °C, and recordings were performed during constant perfusion with temperature measured using a thermistor (TA-29, Warner Instruments) located close to the cell. The thermistor was connected to the digitizer via a temperature controller (TC-324B, Warner Instruments) (Zhang et al., 2018). Oocytes injected with the various constructs were only studied if un-injected oocytes from the same batch contained a low density of endogenous temperature-sensitive currents at negative membrane voltages. For delivering cold stimuli, the external solution was passed through glass capillary spirals that were immersed in an ice bath. The mean log(I) vs T relations were calculated by automatically binning each of the recordings in temperature intervals of 2 °C, and averaging a single data point within each interval per cell. All data analysis was carried out using Igor Pro 6.3 (Wavemetrics, Portland, OR).

### Reagents and chemicals

All reagents were obtained from Sigma-Aldrich (St. Louis, MO), unless indicated otherwise. Resiniferatoxin (RTx, Tocris Bioscience, Bristol, UK) and capsaicin stock solutions were made in ethanol. 2-APB stock solutions were made fresh every day in DMSO. Double knot toxin (DkTx) was synthetized in the laboratory as described previously(Bae et al., 2012).

## Acknowledgments

We thank Shai Silberberg, Gilman Toombes and members of the Swartz lab for helpful discussions. This work was supported by the Intramural Research Programs of the NINDS, NIH (to K.J.S.), by an NINDS Competitive Postdoctoral Fellowship and K99 Career Development Award (to A.J.O).

## Author contributions

F.Z. performed experiments and all authors contributed to the design of the study and the writing of the manuscript.

---

**Figure 1 Supplement 1.**
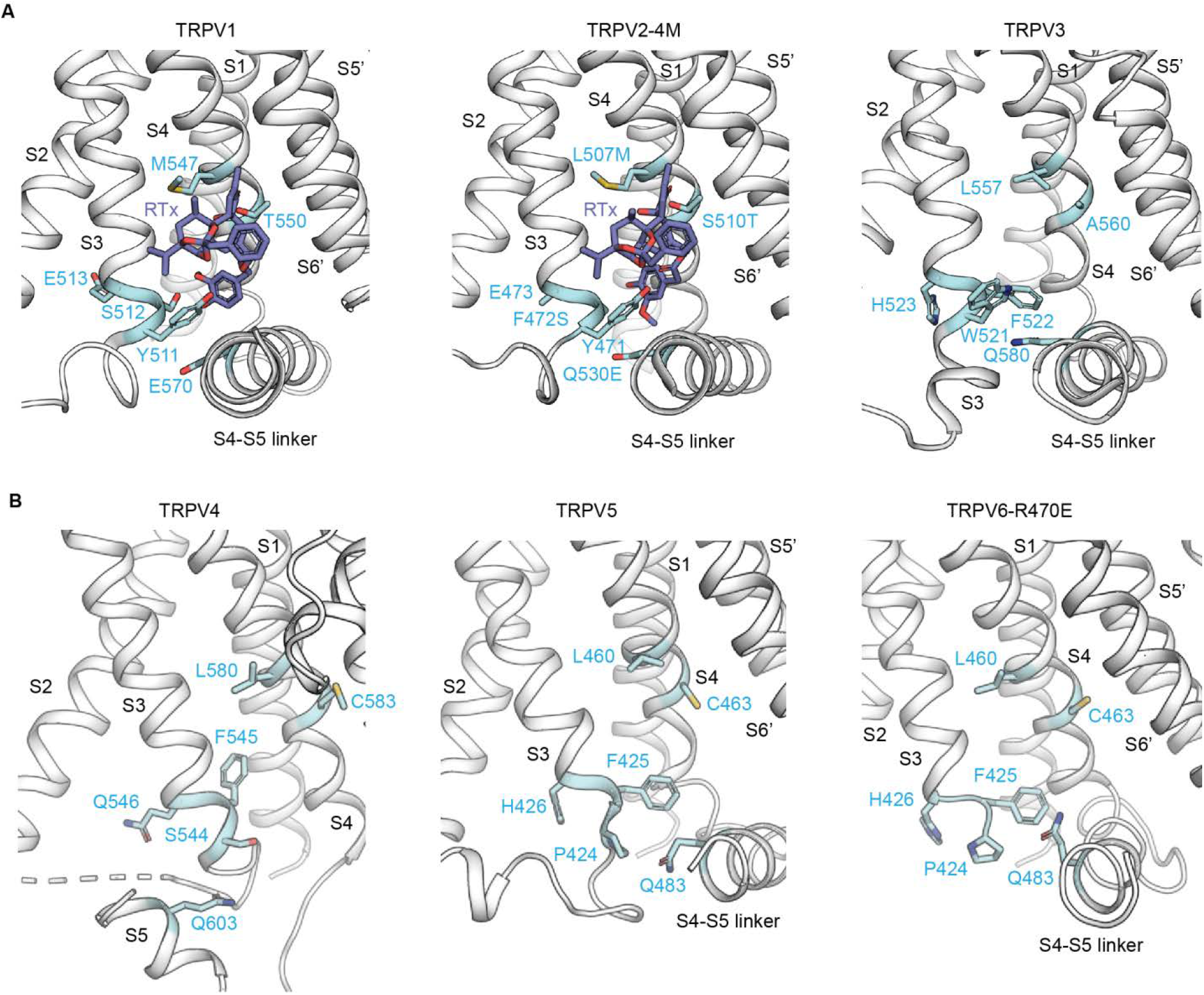
The vanilloid binding pocket in the TRPV family. **(A)** Cartoon representation of the vanilloid-binding pocket formed by the S1-S4 helices of one subunit and the pore domain of the adjacent subunit (S5’ and S6’) depicted for rat TRPV1 (in complex with RTx and DkTx, PDB: 5IRX) (Gao et al., 2016), rabbit TRPV2-4M (in complex with RTx, PDB: 6BWJ) (Zubcevic et al., 2018b) and mouse TRPV3 (in complex with 2-APB, together with the Y564A mutation, PDB: 6DVZ) (Singh et al., 2018). The residues corresponding to the 6M mutations are depicted in light blue. RTx is shown in dark blue. Structures are shown in equivalent orientations based on structural alignments obtained in Pymol with the cealign command. **(B)** Cartoon representation of the regions corresponding to the vanilloid-binding pocket for zebrafish TRPV4 (PDB: 6BBJ) (Deng et al., 2018), rabbit TRPV5 (in complex with econazole, PDB: 6B5V) (Hughes et al., 2018) and human TRPV6 (R470E mutant, PDB: 6BOA) (McGoldrick et al., 2018), aligned as in (A). The residues equivalent to the 6M mutations are shown as sticks in light blue. The S6’ was omitted from the TRPV4 figure to allow clearer visualization of the pocket.

**Figure 3 Supplement 1.**
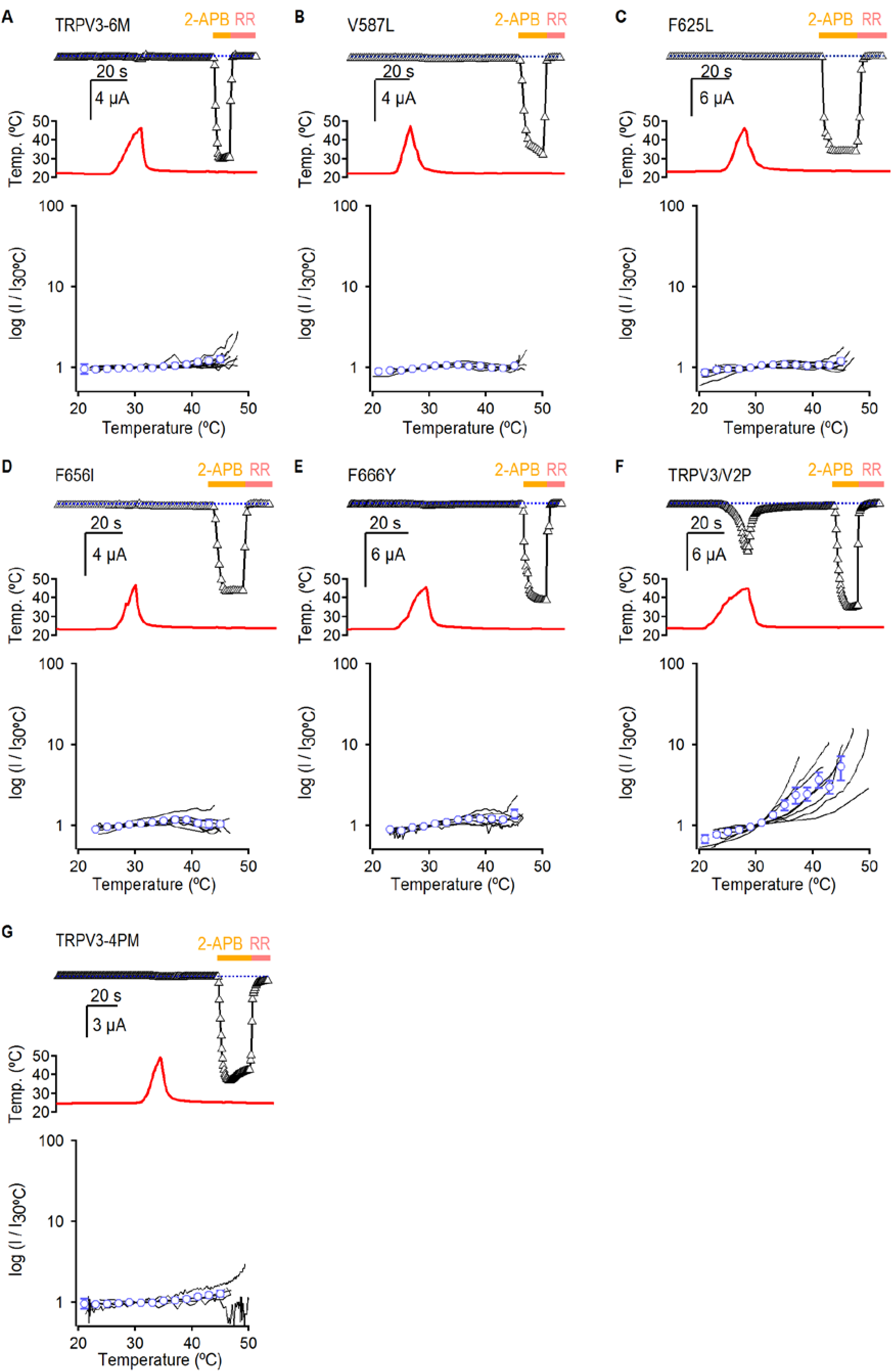
Temperature-activation of TRPV3 constructs. (A-G) The top panels show representative time-courses obtained as in Fig. 3, depicting the response to temperature and 2-APB of TRPV3 constructs. The dotted lines indicate the zero-current level. The middle panels show the temperature measured during each experiment depicted on the top panels. The bottom panels show log(current)-temperature relations at -60 mV obtained from experiments as in the upper panels. Individual cells are shown as black curves, with currents normalized to their amplitude at 30 °C, and the mean ± S.E.M are shown as open blue circles (n=5-8).

**Figure 4 Supplement 1.**
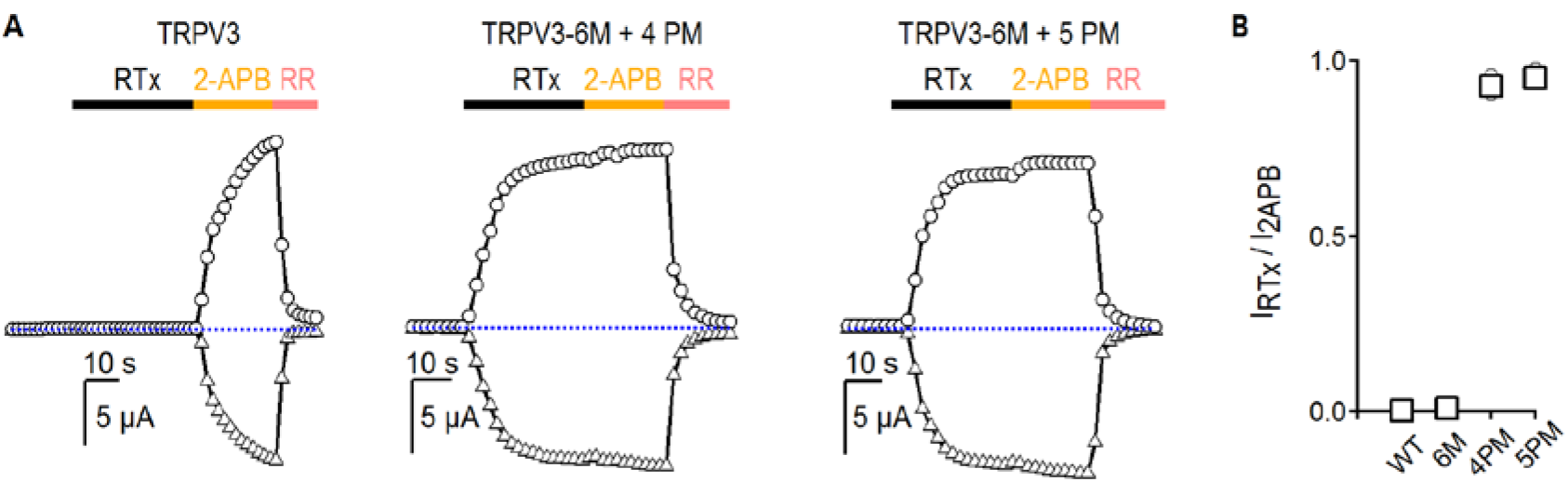
Relative maximal activation by RTx and 2-APB for TRPV3 constructs. **(A)** Representative time courses (−60, triangles; +60 mV, circles) of activation by RTx (100 nM) and 2-APB (3 mM), followed by block with RR (50 μM). The dotted horizontal lines indicate the zero-current level. **(B)** Summary of the sensitivity of TRPV3 constructs at +60 mV to RTx relative to 2-APB from experiments as in (A). Values for individual oocytes are shown as open grey circles and the mean ± S.E.M. as bars (n = 3-4).

**Figure 5 Supplement 1.**
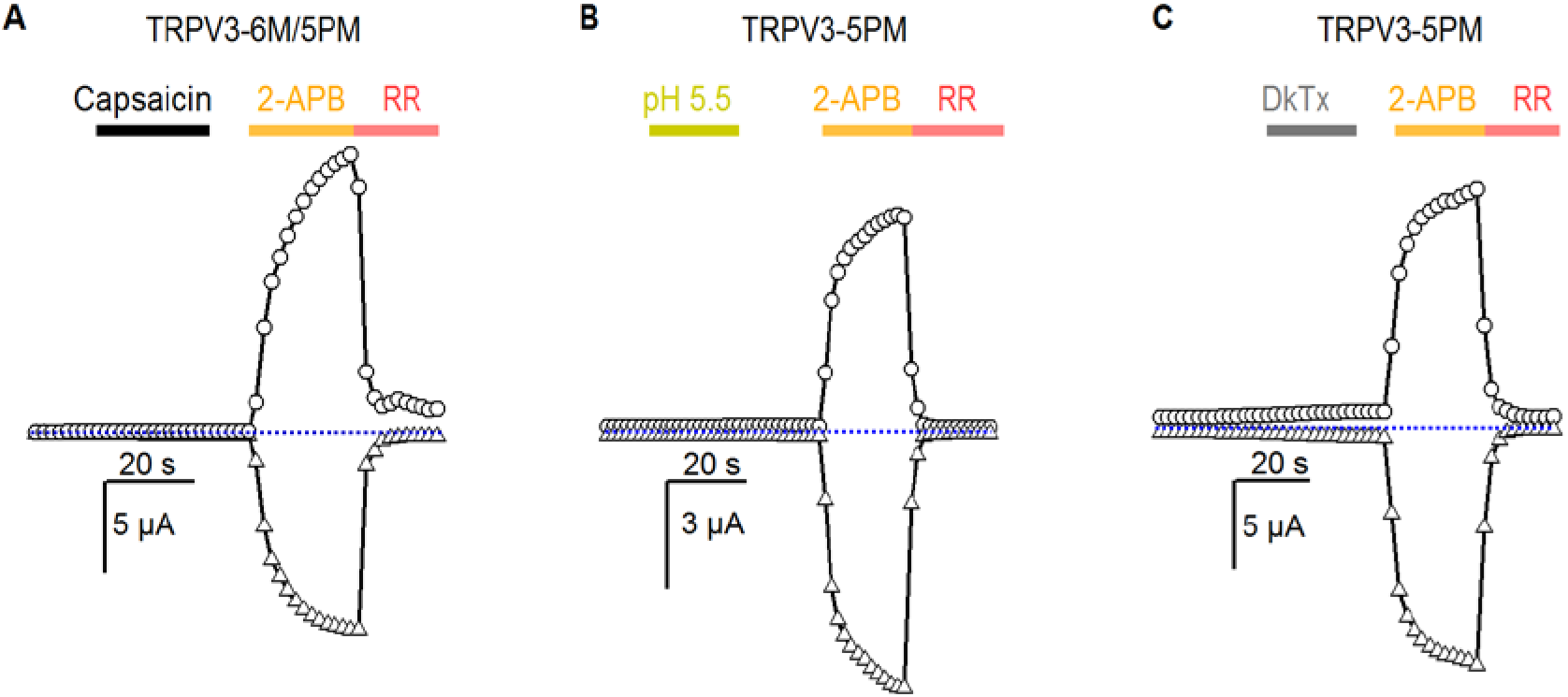
Pore mutants of TRPV3 do not enable sensitivity to other TRPV1-specific agonists. (A) Representative current time courses at +60 mV (circles) and -60 mV (triangles) showing that TRPV3-6M/5PM channels cannot be activated by a high concentration of capsaicin (20 μM). The dotted line denotes the zero-current level. **(B, C)** Representative time courses at +60 mV (circles) and -60 mV (triangles) showing that TRPV3-5PM channels cannot be activated by (B) low pH or (C) the double-knot toxin (1 μM, DkTx). 2-APB was used at 3 mM, and RR at 50 μM.

